# Fluorescent sensors for activity and regulation of the nitrate transceptor CHL1/NRT1.1 and oligopeptide transporters

**DOI:** 10.1101/002741

**Authors:** Cheng-Hsun Ho, Wolf B. Frommer

## Abstract

To monitor nitrate and peptide transport activity *in vivo*, we converted the dual-affinity nitrate transceptor CHL1/NRT1.1/NPF6.3 and four related oligopeptide transporters PTR1, 2, 4 and 5 into fluorescence activity sensors (NiTrac1, PepTrac). Substrate addition to yeast expressing transporter fusions with yellow fluorescent protein and mCerulean triggered substrate-dependent donor quenching or resonance energy transfer. Fluorescence changes were nitrate/peptide-specific, respectively. Like CHL1, NiTrac1 had biphasic kinetics. Mutation of T101A eliminated high-affinity transport and blocked the fluorescence response to low nitrate. NiTrac was used for characterizing side chains considered important for substrate interaction, proton coupling, and regulation. We observed a striking correlation between transport activity and sensor output. Coexpression of NiTrac with known calcineurin-like proteins (CBL1, 9; CIPK23) and candidates identified in an interactome screen (CBL1, KT2, WNKinase 8) blocked NiTrac1 responses, demonstrating the suitability for *in vivo* analysis of activity and regulation. The new technology is applicable in plant and medical research.

## Introduction

Quantitatively, nitrogen is the single most limiting nutrient for plants. Thus, not surprisingly, maximal crop yield depends critically on nitrogen fertilizer inputs. Current practices require production of ~1.5 10^7^ tons of N-fertilizer *per annum,* consuming ~1% of the world’s annual energy production. Plants absorb only a fraction of the fertilizer applied to the field, leading to leaching into groundwater, polluting the environment and damaging human health. Improvements in nitrogen use efficiency of crops are urgently required, however, while potential targets including uptake transporters and metabolic enzymes have been identified, successful improvements in N-efficiency are rare {McAllister, 2012 #47168;Jiang, 2012 #47174;Xu, 2012 #47195;Schroeder, 2013 #38704}. Overexpression of an alanine amino transferase or the transporter *OsPTR9* are two of the few examples of improved nitrogen use efficiency {Shrawat, 2008 #47212;Fang, 2013 #47196}. Ammonium, nitrate, amino acids and di- and tripeptides serve as the major forms of inorganic and organic nitrogen for plants. Uptake occurs predominantly from the soil/rhizosphere into roots, although aerial parts of the plant are also capable of absorbing nitrogen {McAllister, 2012 #47168}. Nitrogen availability and distribution in soil vary both spatially and temporally. Inorganic nitrogen uptake is complex and involves multiple ammonium and nitrate uptake systems, typically grouped into low-affinity/high capacity and high-affinity/low capacity systems {Siddiqi, 1990 #4490;Wang, 1994 #46939;von Wirén, 1997 #271}. Their relative activity is influenced by both exogenous and endogenous factors. The exact sites of uptake of the various forms of nitrogen along the length of the root, the cells that are directly involved and *in vivo* regulation are not well understood. Also the exact intercellular path towards the stele is not experimentally proven. The reasons for this lack of knowledge lie in the fact that nitrogen transport is difficult to measure. Some studies rely on the analysis of the depletion of the medium, others use stable isotopes, or the ^13^N-isotope, which has a short half-life time of ~10 minutes and requires access to a suitable supply source {Clarkson, 1996 #47227;Wang, 1993 #46942}. Most of these techniques lack spatial resolution, i.e. information on which cell layers and which root zones absorb the nutrient. Electrophysiological assays can provide spatial information, however, they are mostly used at accessible surfaces. Spatial information has been provided in a few studies by methods such as vibrating electrodes {Henriksen, 1990 #47191;Henriksen, 1992 #47192}, positron-emitting tracer imaging systems {Kiyomiya, 2001 #4957;Matsunami, 1999 #47226} or Secondary Ion Mass Spectrometers {Clode, 2009 #47224}. We also know little about differences in the distribution of the nitrogen forms in different root cell types or zones and with respect to cellular compartmentation. Classical approaches average total ion/metabolite levels over all cells in the sample, e.g. in whole roots. Nitrate levels differ dramatically between root cell types {Karley, 2000 #5731;Zhen, 1991 #47236}. Recently a GFP-labeled protoplast-sorting platform was used to compare metabolomes of individual cell types in roots {Moussaieff, 2013 #47231}. This study found that levels of small oligopeptides were comparatively higher in the epidermis and endodermis compared to other root cell types. Compartmental analyses indicated that the nitrate concentration of root vacuoles is ~10-fold higher compared to the cytosol {Henriksen, 1990 #47236}.

Transporters are placed in strategic positions to control which and how much of a specific nitrogen form can enter a given cell at a given point of time. The progress in identifying transporter genes provided a new handle for addressing the mechanisms and the spatial and temporal regulation of nitrogen acquisition from a new level of detail. Three major families of transporters for inorganic nitrogen uptake (and distribution) have been identified: the NPF/POT nitrate transporter family {Leran, 2013 #47172}, NRT2 nitrate transporters {Kotur, 2012 #47180}, and the ammonium transporters of the AMT/MEP/Rh family {Andrade, 2007 #46873;von Wirén, 2000 #1807}. In addition to their role in nitrate uptake, members of the NPF/POT {Leran, 2013 #47172} family play important roles also in the transport of histidine, dicarboxylates, oligopeptides, glucosinolates and surprisingly at least three major plant hormones: auxin, ABA and gibberellin {Boursiac, 2013 #46967;Krouk, 2010 #47183;Kanno, 2012 #47030}. Genes are valuable tools for exploring physiological functions. Analysis of RNA levels allows us to study gene regulation {Gazzarrini, 1999 #4798}, e.g. transcriptional GUS-fusions for determining organ and cell type specific expression, translational GFP-fusions for subcellular localization. Both classical and novel methods, including cell-specific transcriptional profiles and ‘translatomes’ provide us with new insights into differences in expression of transporters in roots {Brady, 2007 #37928;Mustroph, 2009 #46141}. Analysis of cell type-specific expression profiles showed that the majority of changes in nitrate-induced gene expression are cell-specific {Gifford, 2008 #47229}. Expression and purification of the proteins followed by reconstitution in vesicles or expression in heterologous systems to interrogate biochemical properties include K_m_ and transport mechanisms. The genes can be used as a basis for structure function studies {Loqué, 2007 #46838} and to obtain crystal structures {Andrade, 2005 #46878;Doki, 2013 #47232}. We can use the genes to identify interacting proteins {Lalonde, 2010 #46051}. Importantly, the availability of genes enables us to generate specific mutants {Wang, 2009 #47239;Yuan, 2007 #46894}, which provide insights into their physiological roles. However, even with this massive amount of detailed data, the key information is missing, namely the information on the activity state of a given protein *in vivo. In vivo* activity depends mainly on two additional parameters beyond protein abundance at a given membrane: the local concentration of the substrate/s, the status of the cell (e.g., the membrane potential and local pH as key determinants for ion transporter activity) and the status of cellular regulatory networks required for activity of the protein in question. Again genes can help us to find regulators and study the effect of mutations on nitrogen acquisition, but ultimately, we need to be able to quantify the activity of the transporters in individual cells *in vivo*.

Nitrogen uptake is controlled by many factors, such as nitrogen level, energy status of the plant, assimilation status of imported nitrogen, N-demand, and involved mobile signals between shoots and roots as well as between different parts of the root system {von Wirén, 1997 #271}. Nitrate transporters are regulated through phosphorylation, mediated by calcium-dependent calcineurin-like kinases (Calcineurin B-like, CBL and CBL-interacting protein kinase, CIPK) {Ho, 2009 #6414;Hu, 2009 #6258;Wang, 2009 #47239}. Major breakthroughs were findings that indicate both members of the AMT and NPF/POT family function as transporters and receptors (transceptors) {Rubio-Texeira, 2010 #46710;Ho, 2009 #6414;Lima, 2010 #46884}. However, despite broad progress, at present, we have only a limited understanding of signaling pathways that control nitrogen acquisition.

It is important to develop tools for monitoring the activity of individual transporters in specific locations in individual cells of plant roots in a minimally invasive manner. A minimally invasive tool that has proven valuable for monitoring ions and metabolite levels with high spatial and temporal resolution are genetically encoded fluorescent nanosensors {Okumoto, 2012 #46924}. These sensors rely on substrate-binding-dependent conformational rearrangements in a sensory domain. The rearrangements are reported by changes in Förster Resonance Energy Transfer (FRET) efficiency between two fluorescent proteins, which act as FRET donor and acceptor due to spectral overlap. Sensors for glucose, sucrose and zinc have successfully been used in Arabidopsis to monitor steady state levels as well as accumulation and elimination under both static and dynamic conditions where roots were exposed to pulses of the respective analytes {Lanquar, 2013 #47222;Deuschle, 2006 #5819;Chaudhuri, 2011 #38398;Chaudhuri, 2008 #7389;Okumoto, 2008 #6844}.

The recent progress in obtaining crystal structures for transporters, and more importantly the availability of transporter structures in multiple configurations, has provided insights into the conformational rearrangements occurring during the transport cycle {Henderson, 2013 #47169;Doki, 2013 #47232;Guettou, 2013 #47175;Madej, 2013 #38678}. Biochemical and structural analyses have shown that many transporters undergo conformational changes during the transport cycle {Shimamura, 2010 #38424;Jiang, 2012 #38425;Krishnamurthy, 2012 #38423}. Important in this context is that such rearrangements have been observed for many members of the MFS superfamily; including members of the NPF/POT family {Doki, 2013 #47232}. We therefore hypothesized that it should be possible to ‘record’ the conformational rearrangements that occur during the transport cycle in a similar manner as used for the engineering of the FRET sensors. The first prototype for transport activity sensors, named AmTrac, uses ammonium transporters as sensory domains for engineering transport activity sensors by inserting a circularly-permutated EGFP (cpEGFP) into a conformation-sensitive position of an ammonium transporter {De Michele, 2013 #38714}. Addition of ammonium to yeast cells expressing the AmTrac sensor trigger concentration-dependent and reversible changes in fluorescence intensity {De Michele, 2013 #38714}. Whether this approach is transferable to other family proteins in different species remained to be shown. To create nitrate and peptide transport activity sensors, we fused CHL1 and four PTRs to fluorescent protein pairs, expressed the fusions in yeast and tested their response to substrate addition (named NiTrac for <u>ni</u>trate <u>tr</u>ansport <u>ac</u>tivity and PepTrac for <u>pep</u>tide transport <u>ac</u>tivity). The five sensors responded to addition of nitrate or peptides, respectively. The kinetics of the NiTrac1 sensor response was strikingly similar to the transport kinetics of the native CHL1; the response was specific and reversible. The new sensors were used to study structure/function relationships, to correlate effects of mutations in CHL1 and NiTrac1 on activity and sensor responses, and to observe the effect of potential regulators on the conformation of the transporter. The successful use of the sensors in yeast indicates that these new tools can be used for *in planta* analyses.

## Results

### Engineering of a nitrate transport activity sensor

It is likely that the nitrate transceptor CHL1 undergoes conformational rearrangements during its transport cycle. To measure substrate-dependent conformational rearrangements, CHL1 was sandwiched between a yellow acceptor (Aphrodite) and cyan donor fluorophore (mCerulean) {Rizzo, 2006 #47257}; Fig. 1A). This chimera, named NiTrac1, was expressed in yeast, followed by spectral analysis of yeast cultures in a spectrofluorimeter (Fig. 1B). The fluorophores were in Förster distance, as evidence by significant resonance energy transfer. If conformational rearrangements were induced by substrate addition, one might expect a change in the energy transfer rate. To our surprise, and in contrast to typical FRET sensors (e.g. glucose or glutamate {Fehr, 2003 #5689;Okumoto, 2005 #6093}), we observed an overall reduction in the emission intensities of both donor and acceptor, but no obvious change in FRET efficiency. Cyan FPs are typically robust compare to the yellow variants; specifically, they are less sensitive to pH changes or other ions compared to yellow variants (here Venus encoded by codon-modified Aphrodite gene sequence {Deuschle, 2006 #5819}). However, Aphrodite emission was unaffected by nitrate when excited directly (Fig. 1B, inset), indicating that external nitrate triggers donor quenching in the cytosol. The sensor response can be expressed as a ratio change between the emission intensity of the sensor at CFP excitation relative to YFP emission obtained from acceptor excitation. As one may have expected, the nitrate analog chlorate that lead to the naming of CHL1 (chlorate resistance of the *chl1* mutant) {Tsay, 1993 #26}, also triggered NiTrac quenching (Fig. 1C). The response of NiTrac1 is nitrate- and chlorate- specific; other compounds such as chloride, ammonium, divalent cations and dipeptide had no significant effect (Fig. 1D). When mCerulean was replaced by the corral-derived cyan fluorescent protein mTFP {Ai, 2008 #7408}, we observed FRET, but nitrate addition had no effect on the emission of this variant (Fig. 1E). The mTFP variant, named NiTrac1c (<u>c</u>ontrol) therefore can serve as a control sensor for *in vivo* measurements. Replacement of mCerulean with eCFP, another jellyfish variant, retained the donor-quenching response to nitrate (Fig. 1F). Although we do not understand the mechanism by which nitrate triggers donor quenching, the effect is likely related to a specific property common to mCerulean and eCFP and lacking in mTFP.

**Figure 1.**
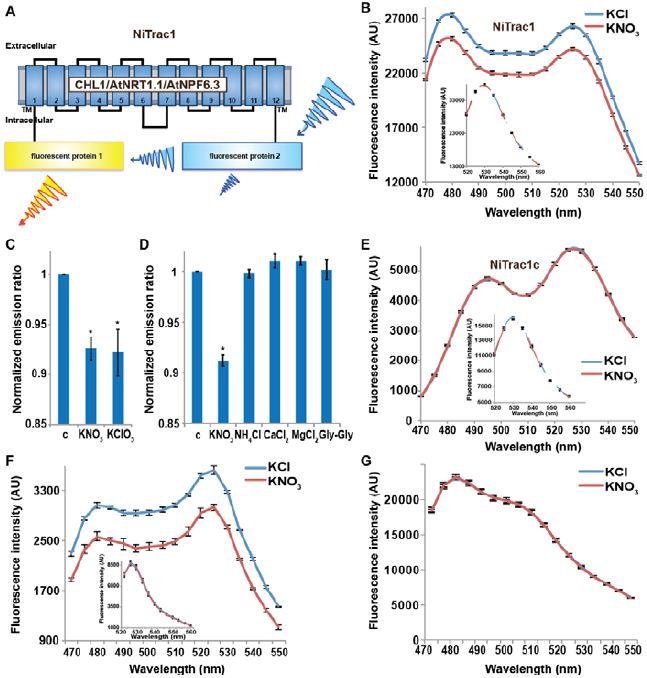
Design and development of NiTrac sensors (**A**). Schematic representation of the NiTrac1 sensor construct. Aphrodite, yellow; mCerulean, light blue; CHL1/NRT1.1/NPF6.3 dark blue; TM, transmembrane domain. (**B**). Emission spectra for NiTrac1 expressed in yeast cells; excitation at 428 nm: addition of 5 mM potassium nitrate (red; control 5 m M KCl - blue) lead to a reduction in fluorescence intensity of donor and acceptor emission, caused by donor quenching. Inset: Emission of Aphrodite in NiTrac1 when excited at 505 nm. Aphrodite emission was unaffected. (**C**). Nitrate and its analog chlorate both trigger quenching at 5 mM concentrations. Nitrate-induced ratio change (peak fluorescence intensity of Aphrodite excited at 505 nm over emission spectrum at 485nm obtained with excitation at 428 nm). Data are normalized to buffer-treated control c (**D**). Substrate specificity: Yeast cells expressing NiTrac1 were treated with the indicated compounds at 5 mM concentrations. Only nitrate and chlorate triggered responses that were significantly different from control c (*, p <0.05, *t*-test). Experiment performed as in Fig. 1C. (**E**). Absence of quenching of NiTrac1 when mCerulean was exchanged for mTFP (excitation at 440 nm). Inset: Emission of Aphrodite in NiTrac1 when excited at 505 nm. (**F**). Donor quenching is retained when mCerulean is exchanged for eCFP in NiTrac1 in response to addition of 5 mM potassium nitrate (red; control 5 m M KCl, blue; excitation at 428 nm). Inset: Emission of Aphrodite in NiTrac1 when excited at 505 nm. (**G**). No detectable effect on the fluorescence properties of nitrate addition to yeast cells coexpresing a cytosolically localized free mCerulean and the CHL1 transceptor. Mean ± SD; n = 3.

### Engineering of four peptide transport activity sensors

It is conceivable that nitrate is taken up by CHL1 into the cytosol where it binds to mCerulean, or eCFP, leading to quenching. However, addition of nitrate to yeast cells expressing CHL1 alone had no effect on the fluorescence of a cytosolically expressed mCerulean (Fig. 1G). One could argue that quenching occurs locally at the exit pore of the transporter directly at the plasma membrane and thus requires tethering of mCerulean to the transporter. To test whether quenching is specifically caused by nitrate, we created similar constructs for the oligopeptide transporters PTR1, 2, 4 and 5 from Arabidopsis {Komarova, 2012 #47264;Tsay, 2007 #38564;Leran, 2013 #47172} (Fig. 2A). These proteins share between 39 and 74% homology with CHL1. PepTrac1, PepTrac2, and PepTrac5 sensors all responded with donor quenching to the addition of 0.5 mM diglycine (Fig. 2B-D). Interestingly, PepTrac4 responded to substrate addition with a ratio change that is consistent with a change in the energy transfer rate rather than donor quenching (Fig. 2E). Further characterization will be necessary to explore the molecular basis of donor quenching and how conformational rearrangements cause donor quenching in NiTrac1 by nitrate or PepTrac1 by peptides, and how they induce resonance energy transfer in PepTrac4.

**Figure 2.**
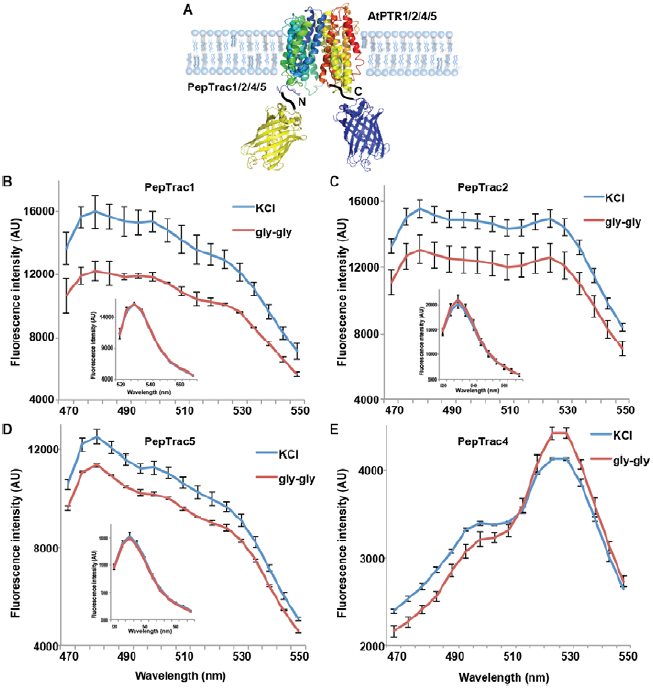
PepTrac sensors. (**A**). Schematic representation of the PepTrac sensor constructs. AtPTR1, 2, 4, and 5 were used for PepTrac sensors creation. Three-dimensional model of AFP-PTRs-mCerulean chimeric protein based on the crystal structure of bacteria peptide transporters (see materials and methods). PTR1 is shown in rainbow cartoon; AFP in yellow; mCerulean in blue. (**B-D**). Donor quenching of PepTrac1, 2, and 5 expressed in yeast in response to addition of 0.5 mM diglycine (red; control 5 mM KCl, blue; excitation at 428 nm). Inset: Emission of Aphrodite in PepTrac1, 2, and 5 when excited at 505 nm. (**E**). FRET ratio change for PepTrac4 (red; control 5 mM KCl, blue; excitation at 428 nm). Mean ± SD; n = 3.

### Biphasic kinetics of the NiTrac1 response

The conformational rearrangements in the sensors could be induced by substrate binding or reflect rearrangements that occur during the transport cycle. Because binding and transport typically have different kinetic constants, we analyzed the response kinetics of NiTrac1. CHL1 is unusual in that it shows biphasic nitrate uptake kinetics (Fig. 3A) {Liu, 2003 #38570}. The observed dual-affinity in oocytes had been attributed to phosphorylation of T101 by endogenous kinase {Liu, 2003 #38570}. The phosphorylation hypothesis would suggest that about half of the transporter molecules are phosphorylated. Interestingly, we observed that the kinetics of the fluorescence response of NiTrac1 in yeast were also biphasic (Fig. 3A). Since it is unlikely that yeast also partially phosphorylates the transporter, the biphasic kinetics are more likely an intrinsic property of the protein. Mutation of T101 to alanine had been shown to eliminate the high-affinity component (Fig. 3B){Liu, 2003 #38570}. Introduction of T101A into NiTrac1 also eliminated the high-affinity component, intimating that NiTrac1 is a transport activity sensor, and that conformational rearrangements during the transport cycle affect mCerulean emission (Fig. 3B). Interestingly, the transport K_m_s of both high and low-affinity phases matched the values obtained for the fluorescence response, supporting the hypothesis that NiTrac measures transport activity. Measurement of the sensor response in individual yeast cells demonstrated rapid nitrate-induced quenching and reversibility of the fluorescence intensity after removal of nitrate (Fig. 3C), indicating that the sensor can be used effectively for *inplanta* analyses.

**Figure 3.**
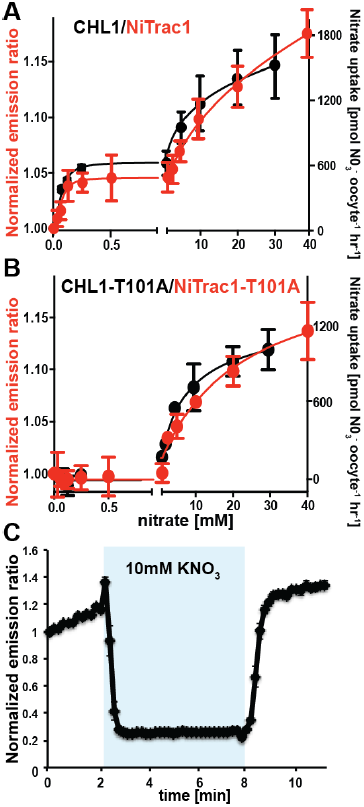
Biphasic kinetics of the NiTrac1 response. (**A**) Biphasic nitrate uptake kinetics of the fluorescence response of NiTrac1 (red) and biphasic nitrate uptake transport kinetics of CHL1/NRT1.1 (Black). (**B**) Monophasic nitrate uptake kinetics of the fluorescence response of NiTrac1-T101A (red) and monophasic low-affinity transport kinetics of CHL1/NRT1.1-T101A (Black, oocyte uptake data from {Liu, 2003 #38570}. The *K*_m_s of NiTrac1 for nitrate are ~75.1±21μM and 3.8±2.6mM; for NiTrac1-T101A is 3.5±3.7mM. Excitation and emission as Fig 1C. The amount of decreased fluorescence intensity by addition of indicated nitrate concentration in Fig 3A and 3B were normalized to water-treated control (0) (mean ± SD; n = 3). (**C**) Analysis of the NiTrac1 response in individual yeast cells trapped in a Cellasic microfluidic plate. Cells were initially perfused with 50 mM MES buffer pH 5.5, followed by a square pulse of 10 mM KNO_3_ in MES buffer for six minutes (blue frame). Data were normalized to the initial value (mean ± SD; n =3).

### Effect of mutations on the NiTrac1 response and NRT1.1 activity

To study the NiTrac mechanism in more detail, and to identify residues important for the transport function of the transporter and sensor, we generated a homology model for CHL1 on the basis of crystal structures of bacterial proton-dependent oligopeptide transporter homologs (see Materials and Methods), and predicted potentially functionally important residues structurally close to the substrate binding pocket from the predicted structure and from sequence alignments. We specifically targeted residues that might be important for substrate specificity, residues involved in proton cotransport, and salt bridges possibly involved in dynamic movements during the transport cycle (Fig. 4A). As one may have expected, different mutants showed different energy transfer ratios, consistent with conformational differences (altered distance and/or orientation of the fluorophores in the absence of substrate; Fig. 4B). Interestingly, we not only observed cases in which donor quenching was lost, but also changes that are consistent with changes in FRET efficiency in response ligand addition, as well as mixtures of donor quenching and change in the FRET efficiency (Fig. 4B). However, without knowledge of the effect of the mutations on transport activity, the data are difficult to interpret. Therefore we introduced the corresponding mutations into CHL1, expressed the mutants in *Xenopus* oocytes and used two-electrode voltage clamp (TEVC, Fig. 5) and ^15^N-uptake (Fig. 6) to measure transport activity. In response to nitrate addition, CHL1 expressing oocytes showed an inward current, consistent with the proposed 2H+/NO_3_^-^ cotransport mechanism. CHL1 contains a highly conserved motif E41-E44-R45 in TM1 predicted to play a role in proton coupling {Doki, 2013 #47232;Solcan, 2012 #47188;Newstead, 2011 #47266;Newstead, 2011 #47267}. Mutations in this motif in the oligopeptide transporters PepT_St_ (from *Streptococcus thermophiles* {Solcan, 2012 #47188}, PepT_So_, PepT_So2_ (both from *Shewanella oneidensis)* {Newstead, 2011 #47267}, and GkPOT (from *Geobacillus kaustophilus)* {Doki, 2013 #47232} typically lost proton-driven transport activity. We therefore tested the role of residues in this motif using NiTrac expressed in yeast and CHL1 expressed on oocytes. Mutation in any of the three residues (E41A, E44A and R45A, TM1) led to a loss of nitrate-induced currents and ^15^N-uptake in both the high- and low-affinity range (0.5/0.25 and 10 mM) (Fig.5, 6). The corresponding mutant of NiTrac1 also lost the sensor response to nitrate addition (Fig.4B, Table 1), indicating that the conserved motif is also used for proton cotransport of nitrate. Interestingly, the mutant was characterized by higher FRET compared to wild type CHL1, indicating that the mutation leads to a conformational change in the protein (Fig. 4B). Structural and functional analyses of the bacterial peptide transporter PepT_St_ had implicated a salt bridge between a conserved K126 in TM4 and E400 in TM10 in peptide recognition and/or structural movements during the transport cycle {Solcan, 2012 #47188}. This lysine is conserved throughout the POT family (K164 of CHL1, TM4) {Doki, 2013 #47232;Solcan, 2012 #47188;Newstead, 2011 #47266;Newstead, 2011 #47267}. Consistent with results from the bacterial PepT_St_ and GkPOT, mutation of K164 to alanine or aspartate in CHL1 completely abolished nitrate uptake in both the high- and low-affinity range (0.25 and 10 mM) (Fig. 6), however, the nitrate-dependent inward currents were retained (Fig. 5). Mutation K136A in GkPOT and K126A in PepT_St_ both abolished completely proton-driven uptake but still had counterflow activities {Doki, 2013 #47232;Solcan, 2012 #47188}. Both CHL1-K164 mutants either function as nitrate-dependent proton channels or have lost selectivity and, consistent with the shift of the reversal potential to more negative values transport other anions such as chloride. NiTrac1-K164A surprisingly showed a different response mode, i.e. upon addition of nitrate the mutant not only showed donor quenching but apparently also a change in FRET efficiency, underlining the exceptional sensitivity of NiTrac1 to effects of mutations on conformation (Fig. 4B). Mutation of the salt bridge acceptor E476A in TM10 of CHL1 led to loss of both the nitrate-induced inward current and ^15^N-uptake (Fig. 5, 6) and NiTrac lost the sensor response to addition of nitrate (Fig. 4B); by contrast, and as one might expect, the conservative mutation E476D had no significant effect on transport properties and sensor response (Fig. 4B, 5, and 6). Alanine substitutions were introduced into corresponding sites predicted to be in the vicinity of the substrate-binding pocket (L49, Q358, and Y388 in TM1, TM7, and TM8, respectively). Consistent with results obtained for the corresponding residue (N342 in TM8) in GKPOT {Doki, 2013 #47232}, Y388A had no detectable effect on the nitrate-induced inward currents, ^15^N-uptake, and sensor response (Fig.4B, 5, and 6), indicating the residue Y388 may not involved in nitrate binding or transport cycle of CHL1. Mutation of L49 in TM1 and Q358 in TM7 of CHL1 to alanine had no significant effect on nitrate-induced inward currents and ^15^N-uptake (Fig. 5, 6), but NiTrac responses were characterized by a mixture of donor quenching and FRET change (Fig. 4B). Based on protein sequence alignments, CHL1 carries an extended cytoplasmic loop connecting the N- and C-terminal six helical bundles. To test the role of this loop, a triple mutant E264A-E266A-K267A was analyzed. The triple mutant lost specifically the low-affinity component nitrate-induced inward current and ^15^N uptake but retained the high-affinity component (Fig. 5, 6), implicating the charged residues in the extended loop in the regulation of nitrate uptake affinity.

**Figure 4.**
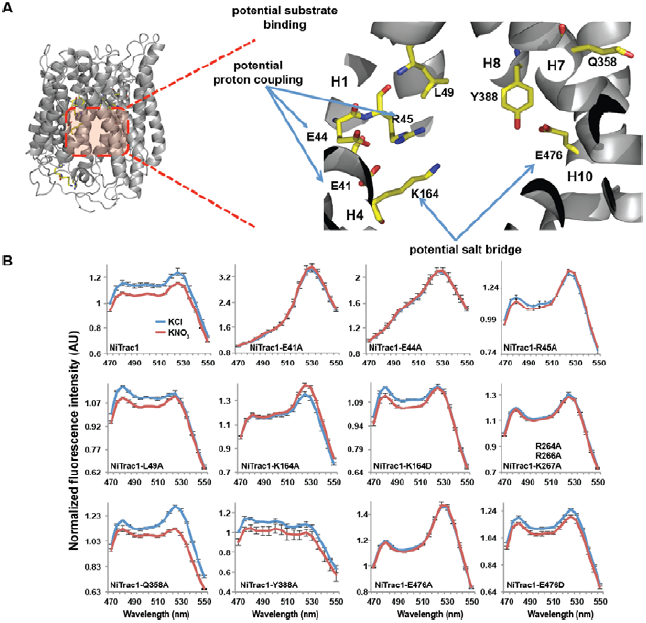
Response of NiTrac1 mutants to nitrate addition. (**A**) Three-dimensional model of CHL1 protein based on the crystal structures of bacteria (see Materials and methods). Red square, potential substrate binding pocket. Left panel, enlarged potential substrate binding pocket. (**B**). Fluorescence response of NiTrac1 mutants expressed in yeast in response to addition of 10 mM potassium nitrate (red; control 10 mM KCl, blue; excitation at 428 nm). To compare the differences in fluorescence intensity between wild type and mutants of CHL1 as well as the differences after addition of nitrate, all data from wild type and mutants were normalized to the intensity of KCl-treated controls at 470nm. Mean ± SD; n = 3.

**Figure 5.**
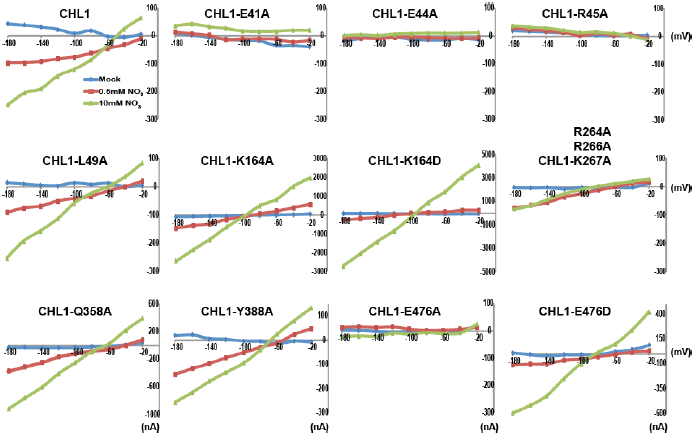
Current and voltage curve of CHL1/NRT1.1 mutants using TEVC. Oocytes were voltage clamped at –40 mV and stepped into a test voltage between −20 and –180 mV for 300 ms, in –20-mV increments. The currents (I) shown here are the difference between the currents flowing at +300 ms in the cRNA-injected CHL1 mutants and water-injected control of the indicated substrates. The curves presented here were recorded from a single oocyte. Similar results were obtained using another two different batches of oocytes.

**Figure 6.**
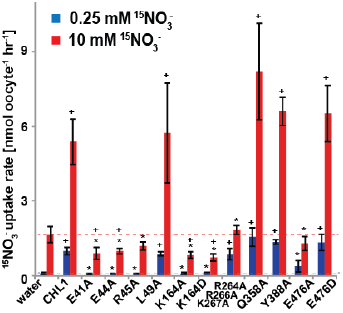
^15^NO_3_^-^ uptake activity of various CHL1 mutants in oocytes. The injected oocytes with various cRNA of CHL1 mutants were incubated with 0.25mM and 10 mM K^15^NO_3_ buffer at pH 5.5 for about 1.5~2 h, and their ^15^N content was determined as described in Methods. The values are the mean ± SD (n = 5~6 for all three experiments). Data are normalized to the 0.25mM treated CHL1-injected oocytes. +, significant difference (p <0.05, *t*-test) compared with water-injected oocytes. An asterisk indicates a significant difference (p <0.05, *t*-test) compared with the CHL1-injected oocytes. Similar results were obtained using another two batches of oocytes.

**Table 1.**
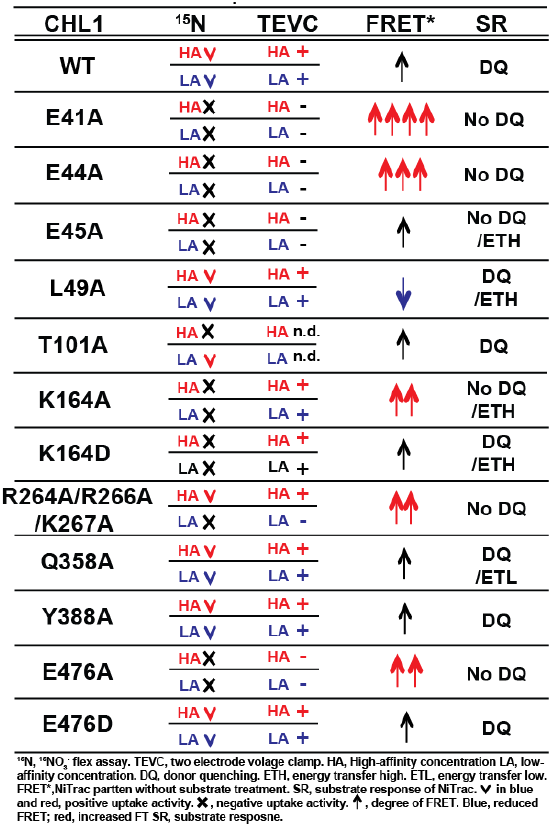
Summary of the nitrate uptake, TEVC, and fluorescence w/wo substrate responses of CHL1 and CHL1 mutants

Similarly, the corresponding NiTrac1mutant also lost the sensor response to high nitrate concentrations (Fig. 4B). Together, these data show that NiTrac1 is a sensitive tool for reporting conformational changes in mutants and further support the hypothesis that NiTrac1 reports activity states of the transporter.

### Effect of regulatory proteins on the NiTrac1 response

The transceptor CHL1 plays important roles in nitrate uptake, transport, sensing, must therefore be subject to regulation of its activity by posttranslational regulation on the one hand, on the other hand CHL1 must interact with intracellular proteins in order to control downstream transcription by signaling pathways. We hypothesized that binding of regulatory proteins or signaling proteins might affect the fluorescence properties of NiTrac1. Therefore, we tested whether coexpression of the known interactor CIPK23, which can phosphorylate CHL1 at T101 in *in vitro* assays, would affect the properties of NiTrac1 (Fig. 7). CIPK23 did not change the energy transfer between the fluorophores in the absence of nitrate, but blocked the fluorescence response of NiTrac1 to nitrate addition (Fig.7B), either by stoichiometric binding or by phosphorylation of T101. The coactivator CBL9, which did not affect CHL1 transport activity on its own but enhanced the CIPK23-mediated phosphorylation of CHL1 {Ho, 2009 #6414}, had no detectable effect on the fluorescence response of NiTrac1 by itself (Fig. 7C). By contrast, CIPK8, which is nitrate inducible in a CHL1-dependent fashion, did not affect the Nitrac1 response. However CBL1 on its own also blocked the Nitrac1 response to nitrate addition (Fig. 7D). The analysis of coexpression of NiTrac1 with combinations of CIPKs and CBLs will require a different approach since episomal expression of three partners likely will create high variability due to copy number variance.

**Figure 7.**
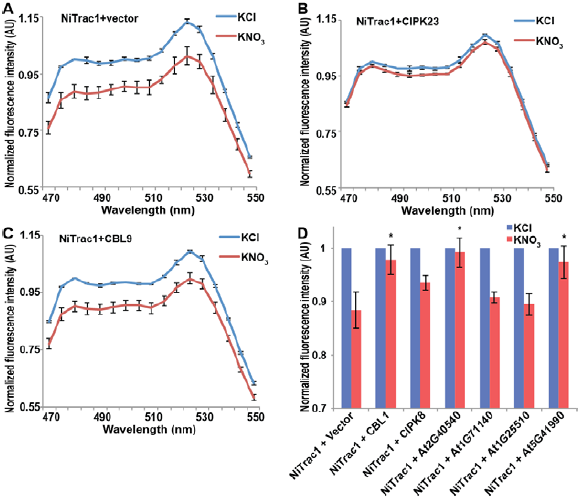
Effects of the fluorescence response of NiTrac1 by interacting proteins. Known interactors or regulators, as CIPK8, CIPK23, CBL1, and CBL9 as well as other interactors identified in a large-scale membrane protein interaction screen were co-expressed with NiTrac1 in yeast cells. (**A**). Donor quenching response of NiTrac1 with vector as control. (**B**) And (**C**). Fluorescence response of NiTrac1 in CIPK23 and CBL9 coexpressing yeast, respectively. The fluorescence response indicates that CIPK23, CBL1, At2g40540, and At5g41990 affect the conformation NiTrac1, whereas no detectable change is observed for CBL9, CIPK8, At1g71140, and At1g25510. (**A-C**). Nitrate-induced ratio change (peak fluorescence intensity of Aphrodite excited at 505 nm over emission spectrum obtained with excitation at 428 nm). Data are normalized to KCl-treated control at 470 nm. An asterisk indicates a significant difference (p <0.05, *t*-test) compared with the KNO_3_-treated control. Mean ± SD; n = 3.

A large-scale interactome screen recently identified novel CHL1 interactors {Jones, 2013 #47204}. To test whether some of these interactors affect NiTrac1 fluorescence, we coexpressed four candidate proteins with NiTrac1 in yeast. While two of the four did not show significant effects on NiTrac1 or the response to nitrate addition, we found that the potassium transporter KT2 and the WNK kinase WNK8 blocked NiTrac1 responses (Fig. 7D). Further experiments will be required to characterize the role of these new interactions; however the results demonstrate the suitability of NiTrac1 for analyzing the effect of known and novel interactors on CHL1 conformation and activity.

### Discussion

To be able to monitor the activity and regulation of individual isoforms of the nitrate and peptide transporter family *in planta* and to study their structure function relationships, we engineered five transporters of the NPF/POT family to report their activity and conformation *in vivo*. The five proteins were fused with yellow and cyan versions of GFP at their N- and C-termini, respectively. When expressed in yeast, the sensors respond to substrate addition either by donor quenching or by a FRET change. The most striking feature of NiTrac is the biphasic response kinetic which matches the dual-affinity transport properties of the protein strikingly well. Based on a predicted structural model and sequence alignments, we mutated select amino acids. Analysis of the fluorescence response of these mutants and comparison with transport assays provides us with insights into the structure-function relationship of the CHL1 nitrate transceptor. The sensor is also used for probing structural rearrangements that occur when NiTrac is coexpressed with putative regulators and interactors, and we discovered new candidate regulatory proteins. The engineering of a suite of nitrate and peptide transport activity sensors complements our recent work in which we developed the prototype for fluorescence-based activity sensors AmTrac and MepTrac by inserting circularly-permutated EGFP into conformation-sensitive positions of ammonium transporters. Addition of ammonium to yeast cells expressing the AmTrac/MepTrac sensors triggered concentration-dependent and reversible changes in fluorescence intensity {De Michele, 2013 #38714}. Together, the engineering of activity sensors through two different approaches - insertion of a fluorophore into a conformationally sensitive site in AMT/MEPs and terminal fusions of a fluorophore pair to the NPF/POT family proteins indicates the potential to transfer the concept to other transporters, receptors, and enzymes. This suite of genetically encoded sensors provides a unique set of tools for observing the activity of individual transporter family members in intact tissue layers of intact plants.

### Sensor output from NiTrac1 and PepTracs

The activity sensors can provide three types of reports: (i) the basic ratio provides information on structure, specifically conformation of the population of sensors which can be compared between for example mutants or in response to coexpression of a regulator; (ii) the intensity of donor or acceptor can be subject to substrate-induced changes that lead to quenching as seen in NiTrac and PepTrac. At present, we do not understand the molecular basis of nitrate-induced donor quenching, which appears to affect mCerulean and CFP, but not mTFP. The fact that three PepTracs show a similar quenching effect when dipeptides are added may indicate that the quenching is caused by a conformational rearrangement in the transporter. A more detailed biophysical characterization may shed light on this unexpected behavior of the sensors. (iii) The change in the emission ratio of the two fluorophores upon substrate addition in PepTrac4 is likely caused by a change in resonance energy transfer as had been observed for small molecule sensors {Okumoto, 2012 #46924}. In certain cases, i.e. NiTrac1 mutants L49A and Q358A (Table 1), we observed a mixture of donor quenching and FRET changes. We thus hypothesize that both NiTracs and all four PepTracs have the potential to report in two different modes, i.e. donor quenching, a FRET change or a combination thereof.

Structural rearrangements triggered by mutations, by binding of a regulator, or by mutations apparently lead to a variety of changes in the fluorescence output. One of the most striking features of NiTrac1 is that it reflects the biphasic kinetics of CHL1 and that even the transport and fluorescence response constants are highly similar. Mutagenesis of T101 to alanine, which had been shown to specifically affect the high-affinity component of nitrate uptake also specifically eliminated the high-affinity response in NiTrac1. These findings strongly supported the notion that NiTrac reports the processes that occur in the transceptor, when it binds and/or transports nitrate. The observations also intimate that the dual-affinity is not caused by partial phosphorylation of CHL1 when expressed in oocytes, as suggested by Tsay’s group {Liu, 2003 #38570}, but more likely represent an intrinsic property of CHL1 since they also occur when NiTrac1 is expressed in yeast. CHL1 also functions as nitrate sensor to regulate transcription of a variety of genes including that of the high-affinity nitrate transporter NRT2 {Ho, 2009 #6414}. Interestingly, the transport and signaling activities of CHL1 can be decoupled by Pro492L in the loop connecting TM10 and 11{Ho, 2009 #6414}. It will thus be interesting to introduce this mutation into NiTrac1 and monitor the effect on the sensor output.

### Using NiTrac1 as a tool for structure function analyses

Taking the advantage of a homology model, we introduced mutations into NiTrac1 and studied the effects on the transport activity by TEVC recording and ^15^N-uptake into oocytes, and compared the effects to fluorescence readouts from the corresponding NiTrac1 mutants. Specifically, we analyzed the role of the putative proton-coupling motif 41ExxER45; the role of charged residues in the extended loop R264R266K267; and residues in the substrate binding pocket as well as a predicted a salt bridge L49, K164, Q358, Y388, and E476 {Doki, 2013 #47232;Solcan, 2012 #47188 Newstead, 2011 #47266;Newstead, 2011 #47267}. We observed three main types of response (Table 1): (i) loss of both nitrate uptake activity and loss of the sensor response in E41A, E44A, R45A, and E476A; (ii) loss of either high- or low-affinity uptake activity and correlated loss of the respective sensor response in T101A and R264A/R266A/K267A; (iii) maintenance of the nitrate uptake activity and sensor response in L49A, Y388A, Q358A, and E476D. Relative to NiTrac1, more than half of the mutants show a change in FRET between the two fluorophores in the absence of substrate addition. Consistent with the role of E41 and E44 in proton-coupling for bacterial peptide transporters, the fluorescence response of E41A, E44A, and R45A NiTrac1 variants support a similar role in the nitrate transporter CHL1. Interestingly, L49A, which did not show detectable differences in transport activity, showed a mixture of donor quenching and FRET change in response to nitrate addition, demonstrating that NiTrac1 is exquisitely sensitive for detecting changes in the overall protein conformation. R45A lost transport activity, but retained a FRET change response after addition of substrate rather than showing a quenching response, indicating an overall conformational change due to binding of nitrate in the absence of a functional transport cycle. Interestingly, mutation of charged residues in the extended cytoplasmic loop of CHL1 (R264A/R266A/K267A) specifically affected the low-affinity component in sensor and uptake response, implicating the loop, potentially through interacting proteins that can tune activity. How T101 phosphorylation, which affects the behavior of CHL1/NiTrac1 at low nitrate levels cooperates with the cytosolic loop, which appears to specifically affect the behavior in high nitrate conditions will be interesting to address in future experiments. Based on our studies, we presume that K164, Q358 and E476 may participate in nitrate binding. Consistent with data from bacterial peptide transporters, E476A lost both sensor response and uptake activity. This conserved residue aspartate likely plays a role in the binding pocket and/or salt bridge formation that is important for the substrate transport cycle. Mutants carrying K164A/D and Q385A mutations were both characterized by significantly increased nitrate-dependent inward currents. It will be interesting to further explore the cause for the increased conductivity with respect to transported ion species. While with the data from the limited number of mutants does not allow us to draw conclusions on the exact molecular nature of the conformational changes, we nevertheless provide the first evidence that activity sensors are highly sensitive and simple tools for probing structure-function relationships in heterologous and homologous systems without the necessity to purify the transporters.

### Effect of CIPK and CBL proteins on the NiTrac1 sensors

The interaction of proteins likely affects the conformation of both partners, either directly or as a consequence of modifications such as phosphorylation. Here we show that activity sensors can be used to probe such interactions with exquisite sensitivity. As a proof of concept, we demonstrate that coexpression of the calcium-dependent kinase CIPK23, which is phosphorylating T101 of CHL1 and thereby inhibiting the low affinity component of CHL1, can block the fluorescence response when coexpressed with NiTrac1. Although typically CIPKs are thought to require a CBL for substrate recognition and derepression of the autoinhibition, CIPK23 had been shown to be able to interact with CHL1 on its own and trigger at least partial phosphorylation of T101 *in vitro* {Ho, 2009 #6414}. Interestingly, although CBL9 had been shown to enhance the CIPK-mediated phosphorylation of T101, we did not observe an effect of coexpression of CBL on NiTrac1. Surprisingly, and despite the high sequence identity between AtCBL9 and AtCBL1 (~89% identity), CBL1 but not CBL9 inhibited the nitrate response of NiTrac1. AtCBL1 and 9 have been shown to regulate a variety of processes including potassium uptake, pollen germination, as well as sugar-, hormone- and ROS-signaling {Xu, 2006 #47038;Cheong, 2007 #47249;Hashimoto, 2011 #47244;Sagi, 2006 #47245;Kimura, 2013 #47250;Drerup, 2013 #47246;Li, 2013 #47251}. Even though CBL1 and 9 have apparent overlapping functions, they can have specific effects, e.g., the AtCBL1-AtCIPK1 complex is involved in ABA-dependent stress responses, while the AtCBL9-AtCIPK1 complex plays roles in ABA-independent stress responses {Drerup, 2013 #47246}. In general, CIPKs depend on their coactivator-CBLs to activate CIPK kinase activity. However, recent studies showed that full-length CIPK23, CIPK16, or CIPK6 alone can activate the AKT1 potassium channel system (Li et al., 2006; Lee et al., 2007; Fujii et al., 2009). Also, AtCBL10 interacts with AKT1 to regulate potassium homeostasis without binding to any AtCIPKs {Ren, 2013 #47034}. The assays deployed here use strong promoters and high copy number plasmids. It will therefore be important to test whether low levels of the kinase are sufficient for inhibiting NiTrac1. It will also be interesting to compare the responses of NiTrac1 when expressed in mutant plants lacking components of the CBL-CIPK machinery.

### WNK kinase and potassium transporter interactions

In addition, we tested whether NiTrac1 can used to monitor conformational rearrangements caused by interacting proteins, specifically we tested interactors detected in a large-scale membrane protein/signaling protein interaction screen {Lalonde, 2010 #46051;Jones, 2013 #47204}. Surprisingly, we found an interaction of CHL1 with the potassium transporter AtKT2/KUP2/SHY3, which plays a role in potassium uptake. Coexpression of KT2 with NiTrac1 lead to a block of the nitrate response. Whether this interaction plays a role in crosstalk between nitrogen and potassium uptake remains to be shown. In addition, we had found an interaction with the *‘no lysine (K) kinase 8’* WNK8. Also WNK8 blocked the nitrate-induced fluorescence response of NiTrac1. WNK8 had been shown to interact specifically with and phosphorylate subunit C of the vacuolar H^+^-ATPase AtVHA-C {Hong-Hermesdorf, 2006 #47279}; as well as with the calcineurin B-like 1 calcium sensor AtCBL1 {Li, 2013 #47280}. It will be interesting to further explore the network between CBL1, WNK8 and CHL1.

Obviously, NiTrac1 is highly sensitive to conformational changes that occur during the transport cycle, effects of mutations and to changes caused by interaction with other proteins. Thus analyses performed with these sensors in plants will have to differentiate between responses caused by substrate and regulatory interactions. The use of controls, e.g. the mTFP sensor, and elimination of FRET by exchanging the acceptor with a non-fretting fluorophore, as well as the use of mutant sensors may be a way to dissect the relative contribution of substrate and protein interactions. These new tools are complementary to the classical tools set including electrophysiology and tracer studies, but has the clear advantage of allowing measurements deep inside plant or animal tissues and organs, domains largely inaccessible to other technologies.

### Outlook

In summary, we developed a set of five sensors that can report the activity of nitrate and peptide transporters *in vivo*. At the same time, such activity sensors prove to be sensitive tools for studying the effect of mutations on the conformation of the transporter or to detect regulatory interactions with other proteins. The next step will be to deploy NiTrac1 and its mutants as well as the PepTracs in Arabidopsis plants to characterize the activity of the transporters and their regulation *in vivo*. The plant peptide transporters are close homologs of the human SLC15 peptide transporters. The SLC15 transporter PepT1 has pathophysiological relevance in processes like intestinal inflammation and inflammatory bowel disease {Ingersoll, 2012 #47274} and it serves as a key transport mechanism for uptake of drugs {Agu, 2011 #47278}. Given the success in engineering five members of the plant transporter family we envisage that the approach can be implemented also for measuring the activity of the human transporters *in situ* and to use such sensors for example for drug screens.

## Materials and Methods

### DNA Constructs

All transporter and sensor constructs were inserted by Gateway LR reactions, into the yeast expression vectors pDRFlip30, 34, 39, and -GW. pDRFlip30 is a vector that sandwiches the insert between an N-terminal Aphrodite t9 (AFPt9) variant {Deuschle, 2006 #5819}, with 9 amino acids truncated of C-terminus, and a C-terminal monomeric Cerulean (mCer){Rizzo, 2006 #47257}. pDRFlip39 sandwiches the inserted polypeptide between an N-terminal enhanced dimer Aphrodite t9 (edAFPt9) and C-terminal fluorescent protein enhanced dimer, 7 amino acids and 9 amino acids truncated of N-terminus and C-terminus of eCyan (t7.ed.eCFPt9), respectively. pDRFlip34 carries an N-terminal AFPt9 and a C-terminal t7.TFP.t9 (t7.TFP.t9){Rizzo, 2006 #47257}. All plasmids contain the f1 replication origin, a GATEWAY™ cassette (attR1-*CmR-ccdB*-attR2), positioned between the pair of fluorescent proteins, the PMA1 promoter fragment, an ADH terminator, and the *URA3* cassette for selection in yeast. Vector construction has been described {Jones, 2013 #44029}. The full length ORF of CHL1, PTR1, PTR2, PTR4, and PTR5 from Arabidopsis and different mutants of NRT1.1 in the TOPO GATEWAY™ Entry vector were used as sensory domains for creating the nitrate sensor NiTrac1, and the peptide sensors PepTrac1, PepTrac2, PepTrac4, PepTrac5. The yeast expression vectors were then created by GATEWAY™ LR reactions between different forms of pTOPO-NRT/PRT and different pDRFlip-GWs, following manufacturer’s instructions. For functional assays in *Xenopus* oocytes, the cDNAs of CHL1 and all mutants of CHL1 were cloned into the oocyte expression vector pOO2-GW {Loqué, 2009 #46882}. Point mutations for studying characterization of CHL1 in oocyte and NiTrac1 in yeast were generated by QuikChange Lightning Site-Directed Mutagenesis Kit (Agilent Technologies). For the coexpression assays with interactors in yeast, putative interactors were inserted, by LR reaction, in the yeast expression vector pDR-XN-GW vector, which was replaced *URA3* with *LEU2* in pDRf1 containing the f1 replication origin, GATEWAY™ cassette (-attR1-CmR-ccdB-attR2), PMA1 promoter fragment, ADH terminator in yeast {Loqué, 2007 #46838}.

### Yeast cultures

The yeast BJ5465 [MATa, *ura3–52, trp1, leu2Δ1, his3Δ200,* pep4::HIS3, prb1Δ1.6R, can1, GAL+] was obtained from the Yeast Genetic Stock Center (University of California, Berkeley, CA). Yeast was transformed using the lithium acetate method {Gietz, 1992 #1477} and transformants were selected on solid YNB (minimal yeast medium without nitrogen; Difco) supplemented with 2% glucose and *-ura/-ura-leu* DropOut medium (Clontech). Single colonies were grown in 5 mL liquid YNB supplemented with 2% glucose and *-ura/-ura-leu* drop out under agitation (230 rpm) at 30°C until OD_600nm_ ~ 0.5 was reached. The liquid cultures were subcultured by dilution to OD_600nm_ 0.01 in the same liquid medium and grown at 30^o^C until OD_600nm_ ~ 0.2.

### Fluorimetry

Fresh yeast cultures (OD_600nm_ ~ 0.2) were washed twice in 50 mM MES buffer, pH 5.5, and resuspended to OD_600nm_ ~0.5 in the same MES buffer supplemented with 0.05% agarose to delay cell sedimentation. Fluorescence was measured in a fluorescence plate reader (M1000, TECAN, Austria), in bottom reading mode using a 7.5 nm bandwidth for both excitation and emission {Bermejo, 2010 #6861;Bermejo, 2011. #38314}. Typically, emission spectra were recorded (λ_em_ 470-570 nm). To quantify fluorescence responses of the sensors to substrate addition, 100 μL of substrate (dissolved in MES buffer, pH 5,5 as 500% stock solution) were added to 100 μL of cells in 96-well flat bottom plates (#655101; Greiner, Monroe, NC). Fluorescence from cultures harboring pDRFlip30 (donor: mCER) and 39 (donor: t7.ed.eCFPt9) was measured by excitation at λ_exc_ 428 nm; cell expressing from pDRFlip34 (donor t7.TFP.t9) were excited at λ_exc_ = 440 nm.

Quantitative fluorescence intensity data from individual yeast cells expressing the sensors (Figure 3C) were acquired on an inverted microscope (Leica, Wetzlar, Germany). To be able to record fluorescence intensities in single cells over time, yeast cells were trapped as a single cell layer in a microfluidic perfusion system (Y04C plate, Onyx, Cellasic, Hayward, CA, USA) and perfused with either 50 mM MES buffer, pH 5.5, or buffer supplemented with 10mM KNO_3_ {Bermejo, 2010 #8121;Bermejo, 2011 #8122}. Briefly, imaging was performed on an inverted fluorescence microscope (Leica DMIRE2) with a QuantEM digital camera (Photometrics) and a 40x/NA (numerical aperture) 1.25–0.75 oil-immersion lens (IMM HCX PL Apo CS). Dual-emission intensity ratios were simultaneously recorded using a DualView unit with a Dual CFP/YFP-ET filter set (ET470/24m and ET535/30m; Chroma) and Slidebook 4.0 software (Intelligent Imaging Innovations). Excitation (filter ET430/24x; Chroma) was provided by a Lambda LS light source (Sutter Instruments; 100%lamp output). Images were acquired within the linear detection range of the camera at intervals of 20 s. The exposure time was typically 1000 ms with an EM (electron-multiplying) gain of 3 °**ø** at 10 MHz and an electron multiplying charge coupled device (EMCCD) camera (Evolve, Photometrics, Tucson, AZ, USA). Measurements were taken every 10 sec, with 100 ms exposure time using Slidebook 5.4 image acquisition software (Intelligent Imaging Innovations, Denver, CO, USA). Fluorescence pixel intensity was quantified using Fiji software; single cells were selected and analyzed with the help of the ROI manager tool.

### Structure prediction for CHL1 and PTR1

Protein structure prediction for CHL1 and PTR1 was performed using Phyre {Kelley, 2009 #47252}. Full-length CHL1 (At1g12110) and AtPTR1/NPF8.1 (At3g54140) amino acid sequences were used for the 3D structure prediction on the website. The analysis made use of 4 solved crystal structures of nitrate/peptide homologs (PDB ID: 4iky, 2xut, 4aps, 4lep){Doki, 2013 #47232;Solcan, 2012 #47188;Newstead, 2011 #47266;Newstead, 2011 #47267}. The homologs shared 16-27% identity with CHL1 or PTR1. The predicted potentially functionally important residues were from the predicted structure (3DLigandSite, {Wass, 2010 #47275}) and from sequence alignments. After structural prediction of CHL1, 41-ExxER-45, in TM1, the conserved sequence motif involved in proton cotransport (22-ExxER-26, 21-ExxER-25, 21-ExxER-25, 32-ExxER-36 in PepT_St_, PepT_So_, PepT_So2_, and GtPOT, respectively), putative residues involved in substrate binding pocket L49 in TM1 (Y30, Y29, Y29, and Y40 in PepT_St_, PepT_So_, PepT_So2_, and GtPOT, respectively), Q358 in TM7 (Q289, Q317, Q291, and Q311 in PepT_St_, PepT_So_, PepT_So2_, and GtPOT, respectively), and Y388 in TM8 (N328, N321, and N342 in PepT_St_, PepT_So2_, and GtPOT, respectively), and putative residues of salt bridges K164 in TM4 (K126, K127, K121, and K136 in PepT_St_, PepT_So_, PepT_So2_, and GtPOT, respectively), E476 in TM10 (E400, E419, E402, and E413 in PepT_St_, PepT_So_, PepT_So2_, and GtPOT, respectively), and residues R264/R266/K267 in the lateral helices loop between TM6 and TM7 were selected for mutagenesis.

### Functional expression of CHL1 and respective mutants in *Xenopus* oocytes

TEVC in oocyte was performed essentially as described previously {De Michele, 2013 #38714}. In brief, for *in vitro* transcription, pOO2-CHL1 and respective mutants were linearized with *Mlu*I. Capped cRNA was *in vitro* transcribed by SP6 RNA polymerase using mMESSAGE mMACHINE kits (Ambion, Austin, TX). *Xenopus laevis* oocytes were obtained from lab of Miriam Goodman by surgery manually, or ordered from Ecocyte Bio Science (Austin, TX). The oocytes were injected via the Roboinjector(Multi Channel Systems, Reutlingen, Germany; {Pehl, 2004 #38504;Lemaire, 2004 #45382}) with distilled water (50 nl as control), or cRNA from CHL1 or CHL1 mutants (50 ng in 50 nl). Cells were kept at 16°C two to four days in ND96 buffer containing 96 mM NaCl, 2 mM KCl, 1.8 mM CaCl_2_, 1 mM MgCl_2_, and 5 mM HEPES, pH 7.4, containing gentamycin (50 μg/μl) before recording experiments. Recordings were typically performed at day three after cRNA injection.

### Electrophysiological measurements in *Xenopus* oocytes

Electrophysiological analyses of injected oocytes were performed as described previously {Huang, 1999 #47253;De Michele, 2013 #38714}. Reaction buffers used recording current (I)-voltage (V) relationships were (i) 230 mM mannitol, 0.3 mM CaCl_2_, and 10 mM HEPES, and (ii) 220 mM mannitol, 0.3 mM CaCl_2_, and 10 mM HEPES at the pH indicated plus 0.5 or 10 mM CsNO_3_. Typical resting potentials were –40 mV. Measurements were recorded by oocytes were voltage clamped at –40 mV and a step protocol was used (-20 to –180 mV for 300 ms, in –20mV increments and measured by the two-electrode voltage-clamp (TEVC) Roboocyte system (Multi Channel Systems){Pehl, 2004 #38504;Lemaire, 2004 #45382}.

### ^15^NO_3_^-^ uptake assays in *Xenopus* oocytes

Nitrate uptake assays were performed using ^15^N-labeled nitrate {Ho, 2009 #6414}, and oocytes injected with CHL1 cRNA were used as positive controls. After two to four days cRNA injection, the oocytes were incubated for 90~120 min in ^15^NO_3_^-^ medium containing 230 mM mannitol, 0.3 mM CaCl_2_, 10 mM HEPES, pH 5.5. Then, oocytes were rinsed five times with ND96 buffer, and individually dried at 80°C for one to two days. ^15^N content was analyzed in an ECS 4010 Elemental Combustion System (Costech Analytical Technologies Inc., Valencia, CA, USA) whose output was connected to a Delta plus Advantage mass spectrometer (Thermo Fisher Scientific Inc., Waltham, MA, USA).

### Statistical analyses

For statistical analyses of ^15^N-nitrate uptake into oocytes (Fig. 6) and the effect of treatments on the fluorescence responses (Fig. 1C and Fig. 7) we used analysis of deviance (ANOVA); factors (sample, treatment) were treated as fixed factors. ANOVAs were performed using the Analysis of Variance (ANOVA) Calculator - One-Way ANOVA from Summary Data (www.danielsoper.com/statcalc). All experiments were performed at least with three biological repeats. The reported values represent mean and standard deviation. Student’s *t*-test was used in Fig. 1, 6, and 7 to determine significance.

## Acknowledgments

We are very grateful to Yi-Fang Tsay for providing constructs and raw data for CHL1 nitrate uptake mediated by CHL1 into oocytes (Fig. 3), and Alexander Jones for providing pDRFLIP vectors. Dr. Ari Kornfeld is gratefully acknowledged for analyzing all ^15^N levels in oocytes shown here. We thank Drs. Newstead and Parker (Oxford University) for providing access to their pre-publication data on the structure of CHL1 and for discussions regarding the transport mechanism. We thank Miriam Goodman (Stanford University) for providing *Xenopus* oocytes. We are grateful to Juan Simón Álamos Urzúa for technical assistance in PepTrac cloning and analysis.

### Additional Information

#### Funding

NSF2010 (MCB-1021677) and NSF 2010 (MCB-1052348) to W.B.F.

#### Author contributions

W.B.F. and C-H.H. conceived and designed the experiments. C-H.H. generated constructs and performed fluorescence and yeast growth experiments as well as oocyte experiments. W.B.F. and C-H.H. analyzed the data. C-H.H. and W.B.F. wrote the manuscript.

